# ZMAP: A single-cell meta-atlas of zebrafish embryonic development reveals a consensus hierarchy of cell identities

**DOI:** 10.64898/2026.03.23.713599

**Authors:** Nicole A. Aponte-Santiago, Yingxin Su, Daniel E. Wagner

## Abstract

Single-cell RNA sequencing (scRNA-seq) efforts have generated large collections of high-resolution cellular atlases of embryonic development, providing unprecedented views of the dynamic gene expression programs that accompany cell fate specification. Alongside parallel efforts in other models, zebrafish has emerged as one of the most extensively profiled vertebrate embryonic systems, with numerous scRNA-seq atlases spanning both embryonic and early larval stages. Despite this progress, cross-study comparisons between datasets remain challenging due to differences in sample processing, mapping, and annotation conventions. Here we present ZMAP (Zebrafish Meta Atlas Project), a harmonized reference integrating 8 published whole-embryo zebrafish scRNA-seq datasets comprising 798,790 cells across 15 developmental time windows. ZMAP unifies component studies through a shared embedding, a standardized marker-gene discovery pipeline, and a hierarchical annotation ontology. Using ZMAP, we inferred “consensus identity programs” – marker gene signatures for each ontology group that were reproducibly detected across studies. To promote broad usage, we provide a Python-based API for automated annotation and retrieval of marker gene sets and reference objects, as well as a web portal that supports interactive 2D and 3D exploration of the UMAP embedding, gene and annotation-level queries, and access to consensus marker resources.

## Background

Recent whole-embryo single-cell RNA sequencing (scRNA-seq) studies spanning several laboratories and technologies have produced extensive, high-resolution maps of zebrafish (Danio rerio) embryonic and larval development (*1–9*). Despite this progress, the collective value of these datasets has not yet been fully realized through systematic integration. Individual studies differ in sampling strategies, technical factors (encapsulation, sequencing, read mapping), and analytic workflows (quality filtering, dimensionality reduction, and clustering). In addition, cell type annotations are often assigned manually by individual laboratories, leading to differences in annotation resolution and descriptive conventions. These inconsistencies complicate cross-study comparisons and limit the ability of the zebrafish community to leverage the full corpus of published scRNA-seq data as a unified reference.

Modern single-cell profiling efforts increasingly leverage the expanding availability of published datasets to construct integrated cross-study meta-atlases (*10*). This strategy enables: (1) assessment of cross-study reproducibility, (2) separation of biological signals from technical covariates, and (3) increased sampling of rare cell states. To extend these advantages to zebrafish, we developed ZMAP (Zebrafish Meta Atlas Project), a harmonized single-cell reference that integrates 8 published whole-embryo zebrafish scRNA-seq datasets. Using this integrated reference, we analyzed cross-study concordance of cell type annotations, identified consensus identity programs, and evaluated ZMAP as a reference for automated annotation and interactive data exploration.

## Results

### Construction of a harmonized zebrafish scRNA-seq reference atlas

We constructed a unified zebrafish single-cell reference atlas by reprocessing and integrating whole-embryo single-cell RNA sequencing data from 8 previously published studies (Fig. 1A) (*1–8*), which were selected based on sequencing depth and overall data quality to ensure reliable cross-study integration. Collectively, these studies span both biological differences (i.e., developmental timepoints; see Fig. 1B) as well as numerous technical differences. To minimize further technical differences due to read processing, raw sequencing reads for each dataset were re-mapped to a common reference genome (see Methods). Library-specific quality control criteria were used to filter low-complexity cell barcodes, cells with elevated mitochondrial transcript fractions, and predicted doublets across 343 individual libraries, yielding a final dataset of over 798,790 high-quality single cells. Summary quality control metrics (e.g., # of mean UMIs and genes detected per cell) varied across studies but were generally consistent with robust single-cell profiling (Fig. 1A).

**Figure 1.**
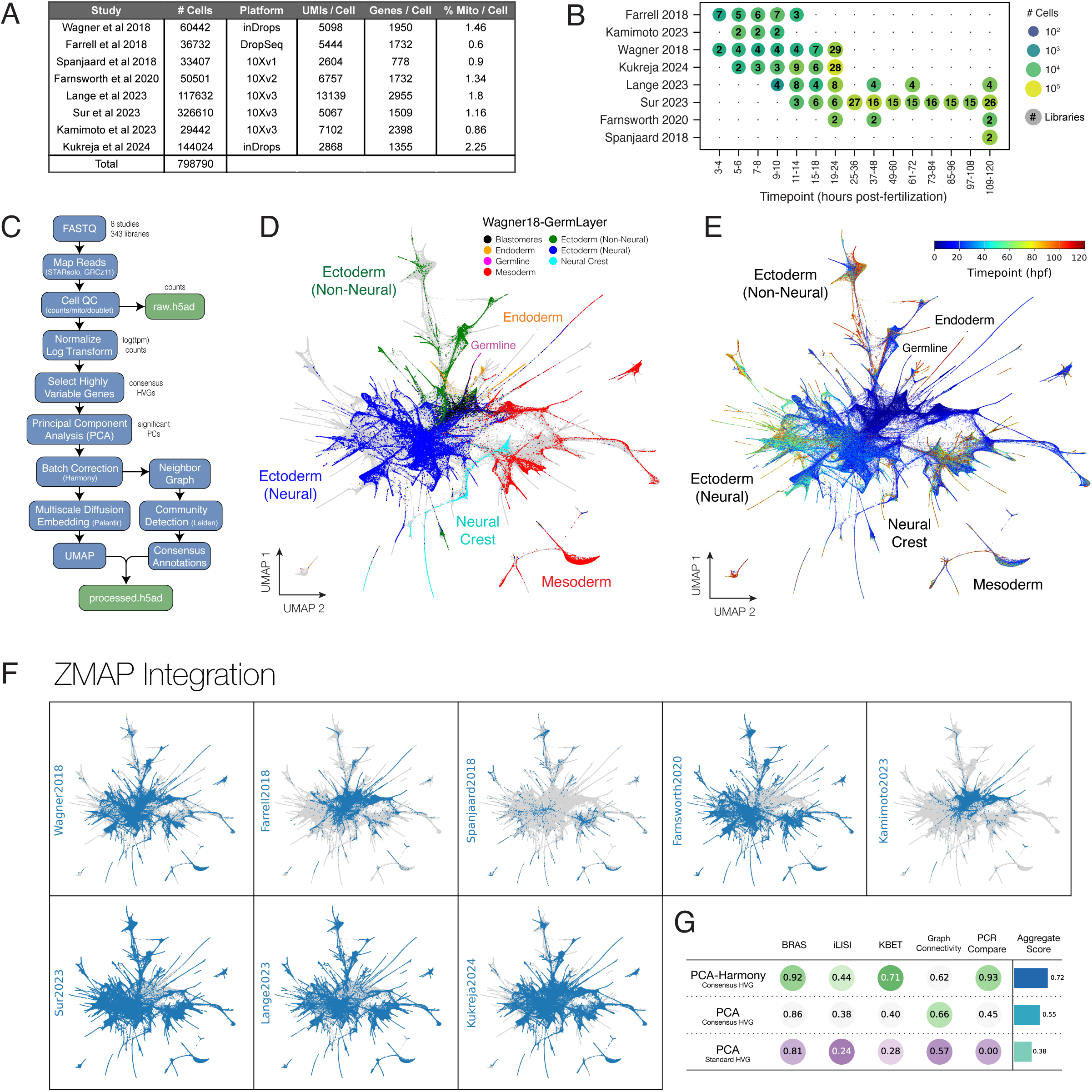
Construction and evaluation of a harmonized zebrafish single-cell reference atlas. (A) Summary table showing # cells retained after quality control, sequencing technology, aggregated quality control (QC) metrics (e.g., detected UMIs and genes per cell, mitochondrial read fraction) for component scRNA-seq studies included in ZMAP. (B) Distribution of cells and libraries across collection time windows (hours post-fertilization; hpf) for each study, visualized as a dot plot, illustrating differences in temporal coverage and sampling density. (C) Data processing and integration workflow, from mapping of raw FASTQ files through cell-level QC, normalization, highly variable gene (HVG) selection, dataset integration, low-dimensional embedding. (D-E) UMAP of the harmonized ZMAP embedding, colored by cell type annotations from Wagner et al. 2018 (D) and by collection timepoint (E). (F) UMAP overlays highlighting study of origin, demonstrating cross-study mixing while maintaining coherent developmental and cell-type structure alongside study-specific differences in temporal coverage. (G) Batch correction and integration performance, using metrics implemented in scIB: Integration Local Inverse Simpson’s Index (iLISI), k-nearest neighbor Batch Effect Test (kBET), graph connectivity, principal component regression (PCR) comparison, Batch Removal Assessment Score (BRAS). Quantifications demonstrate gains following both batch-aware feature (HVG) selection and Harmony-based integration.

ZMAP component libraries derive from overlapping developmental time windows (Fig. 1B), which facilitated an integration workflow including normalization, library/study-aware feature selection, dimensionality reduction, and batch correction (Fig. 1C). Highly variable genes (HVGs) were identified separately for each of the 343 libraries and compared to prioritize features exhibiting reproducible variability across libraries and studies. Principal component analysis (PCA) was performed on this HVG feature set, with the number of significant components estimated using a random permutation-based eigenvalue thresholding approach (see Methods). Residual batch effects associated with library and/or study origin were further corrected using Harmony (*11*), yielding an integrated representation for downstream analysis.

Uniform Manifold Approximation and Projection (UMAP) (*12*) embeddings generated from a Palantir-based multiscale diffusion map (*13*) revealed a continuous developmental manifold structured by both cell identity and time (Fig. 1D-E). Overlay of individual study labels demonstrated substantial cross-study mixing within shared biological states and temporal windows (Fig. 1F).

Batch correction performance was evaluated using scIB metrics (*14*) (Fig. 1G) on a biologically matched subset of 24 hours post-fertilization (hpf) cells, the developmental stage with the greatest study overlap (n = 5 studies; 10,000 cells sampled per study). Integration performance was compared across three procedures: (i) a baseline PCA embedding using “standard” (non-batch aware) HVG selection, (ii) a PCA embedding using batch-aware “consensus” HVG selection, and (iii) a Harmony-corrected embedding derived from (ii). Both Harmony and batch-aware feature selection improved batch integration relative to uncorrected embeddings, as revealed by iLISI, kBET, and principal component regression (PCR) (*14*, *15*). Successful batch correction was also indicated by Batch Removal Assessment Scoring (*14*) (BRAS), a joint measure of cross-study integration and preservation of biological variation.

### A harmonized hierarchical ontology of zebrafish cell type annotations

Most high-quality cell type atlases are constructed through a painstaking process of manual curation using field-specific expertise. Independent curation efforts have thus naturally drifted towards their own styles and conventions in which ontological resolution, decisions between anatomical, positional, and/or cell type-centric identity definitions, and descriptive conventions become fixed. To anchor ZMAP cells into a coherent and unified annotation framework, we utilized a semi-supervised procedure. First, ZMAP cell state neighborhoods were constructed from the Harmony-integrated low-dimensional representation using a k-nearest neighbor (kNN) graph, and graph-based community detection was performed using the Leiden algorithm (*16*). High-resolution clusters (Leiden resolution = 100; 1506 clusters) in which study/batch mixing was well maintained, were used to define the primary units for annotation. Curated “CellType” labels were then assigned to these Leiden clusters by jointly considering the most frequent label from each study (Supplemental Table 1). These CellType labels were subsequently organized into a hierarchical ontology informed by multiple sources of evidence, including lower-resolution Leiden clustering (e.g., resolutions 1, 5, 20), cluster connectivity within the kNN graph, and marker gene expression. The resulting ontology spans five annotation levels: (1) GermLayer, (2) Tissue, (3) CellType (primary curated annotation), (4) CellTypeFine (manually refined subtypes), and (5) the original Leiden algorithm-defined clusters. Together these ontology levels provide a view of cell identity that generalized across the integrated UMAP embedding (Fig. 2A) and across studies (Fig. 2B).

**Figure 2.**
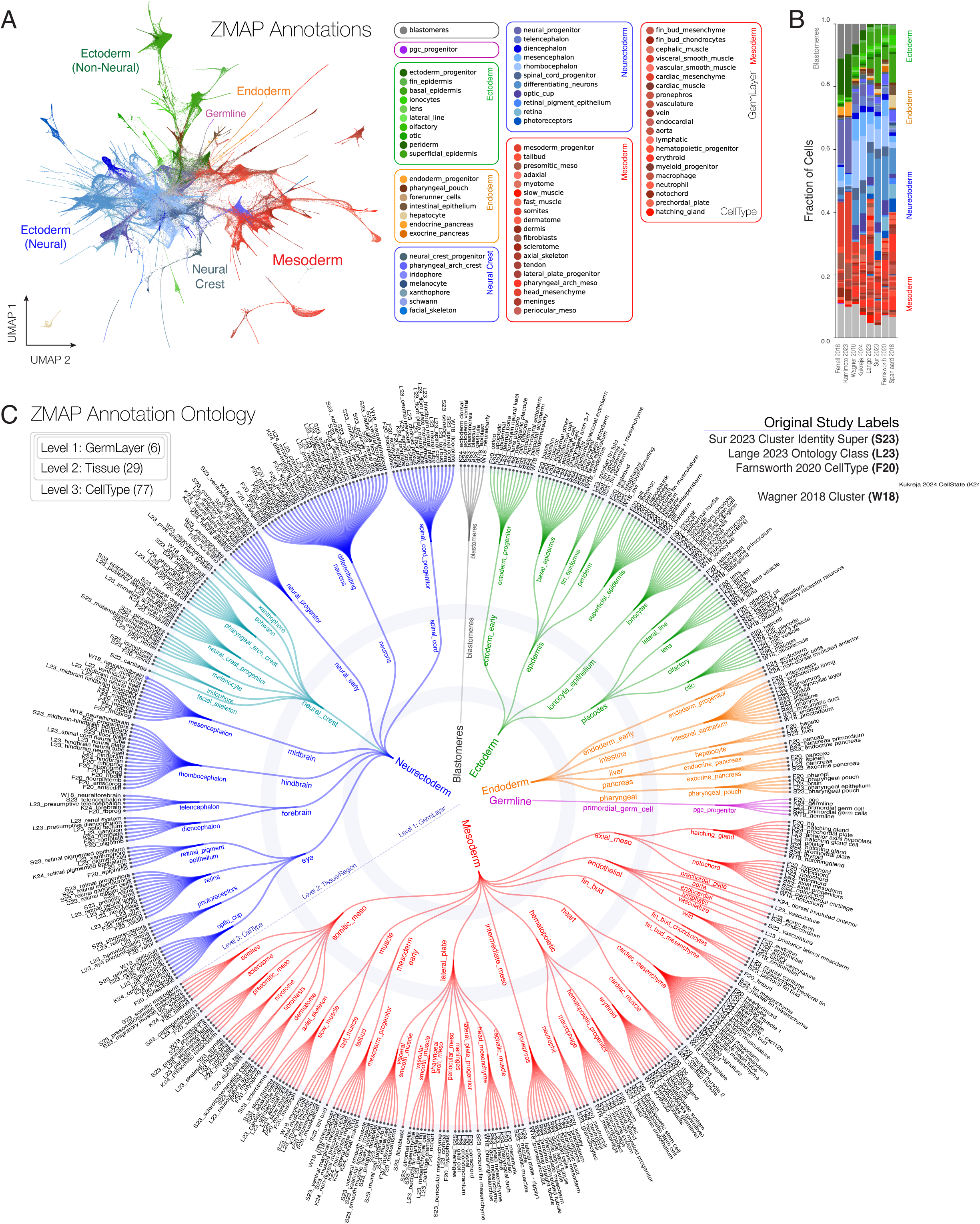
A study-harmonized ontology of cell identity labels. (A) UMAP overlay depicting ZMAP annotation layers. Individual colors correspond to CellType labels while hue groups (boxed) correspond to ZMAP-GermLayer labels. (B) Fractional distributions of the CellType labels in (A) across the 8 component studies. (C) ZMAP ontology relationships across 3 levels: GermLayer (level 1), Tissue (level 2), and CellType (level 3) and associated study-specific labels (level 4), revealing links between semantically similar label sets. Tree edges were determined by maximum connectivity scores between labels, as calculated from the harmonized kNN graph.

To assess semantic convergence, we constructed an ontology tree based on cross-study connectivities between label sets in the integrated kNN graph (Fig. 2C). Connectivities were computed as the sum of single-cell edges linking all possible label pairs across three ZMAP annotation layers (GermLayer, Tissue, CellType) and study-derived annotations. A directed graph was then constructed and rendered as a radial tree, which demonstrated overwhelming cross-study convergence of semantically related groups of labels (Fig. 2C). For example, the ZMAP CellType label for “hatching gland” was associated with study labels: “prechordal plate”, “hatching gland”, “hg”, “polster”, and “anterior axial hypoblast”, synonymous descriptions for derivatives of the dorsal-anterior axial mesoderm. Similar relationships were recovered throughout the ontology. A lookup table, which links each ZMAP annotation to its constituent study labels, is also provided for both programmatic and ad-hoc exploration of these relationships (Supplemental Table 2). The full ZMAP ontology, built from these semantically related groups, thus provides a unified reference vocabulary for subsequent interrogation of zebrafish cell state identity.

### Identification of consensus cell identity programs across studies

To define robust transcriptional programs underlying cell identity, we developed a meta-study differential expression testing pipeline, which assessed gene-level evidence across independent scRNA-seq datasets within ZMAP. Rather than relying on marker genes derived from a single study, this approach identifies genes that reproducibly distinguish the same cell group across multiple experimental contexts, technologies, and laboratories, thus yielding core biological signatures. We refer to these core marker gene sets as “consensus identity genes”. For each annotation group at every level of the ZMAP ontology, candidate identity genes were first detected independently within each contributing study using one-versus-rest non-parametric differential expression testing (see Methods). Only study-group combinations with sufficient representation (≥1% of cells in that group) were considered, ensuring that consensus inference was not driven by rare or weakly sampled populations. Within each eligible study-group pair, candidate identity genes were then filtered to meet thresholds for statistical significance (adjusted p-value ≤ 0.01), effect size (log₂ fold change ≥ 1), and expression prevalence (detected in ≥ 10% of group cells), yielding study-specific marker sets that satisfied both differential expression and specificity criteria. To prioritize genes that best captured each group identity, we combined complementary ranking criteria for several metrics: differential contrast, cell-type specificity, expression prevalence, and cross-study reproducibility (see Methods).

Importantly, this strategy did not enforce a fixed number of markers per group; instead, marker set sizes naturally varied across ontology levels, reflecting differences in transcriptional complexity and internal heterogeneity. Visualization of gene expression patterns across the ontology revealed the structure of cell identity programs (Fig. 3). At coarse ontology levels (e.g., GermLayer) consensus identity genes were broadly expressed and reproducible but differed in their ability to uniformly label groups with high levels of internal state heterogeneity (Fig. 3B). As cluster granularity increased (e.g., Tissue), identity programs became progressively restricted, with some markers exhibiting uniform specificity while others revealed signatures shared between related groups (Fig. 3C). Along selected ZMAP ontology paths, identity gene signatures showed progressive refinement, illustrating nested transcriptional programs across the hierarchy (Fig. 3D). Together, these results demonstrate that consensus identity genes indeed capture stable, biologically meaningful transcriptional programs, which generalize across studies and reflect nested levels of cellular identity. A full table of consensus identity genes associated with each ZMAP ontology group is provided in Supplemental Table 3.

**Figure 3.**
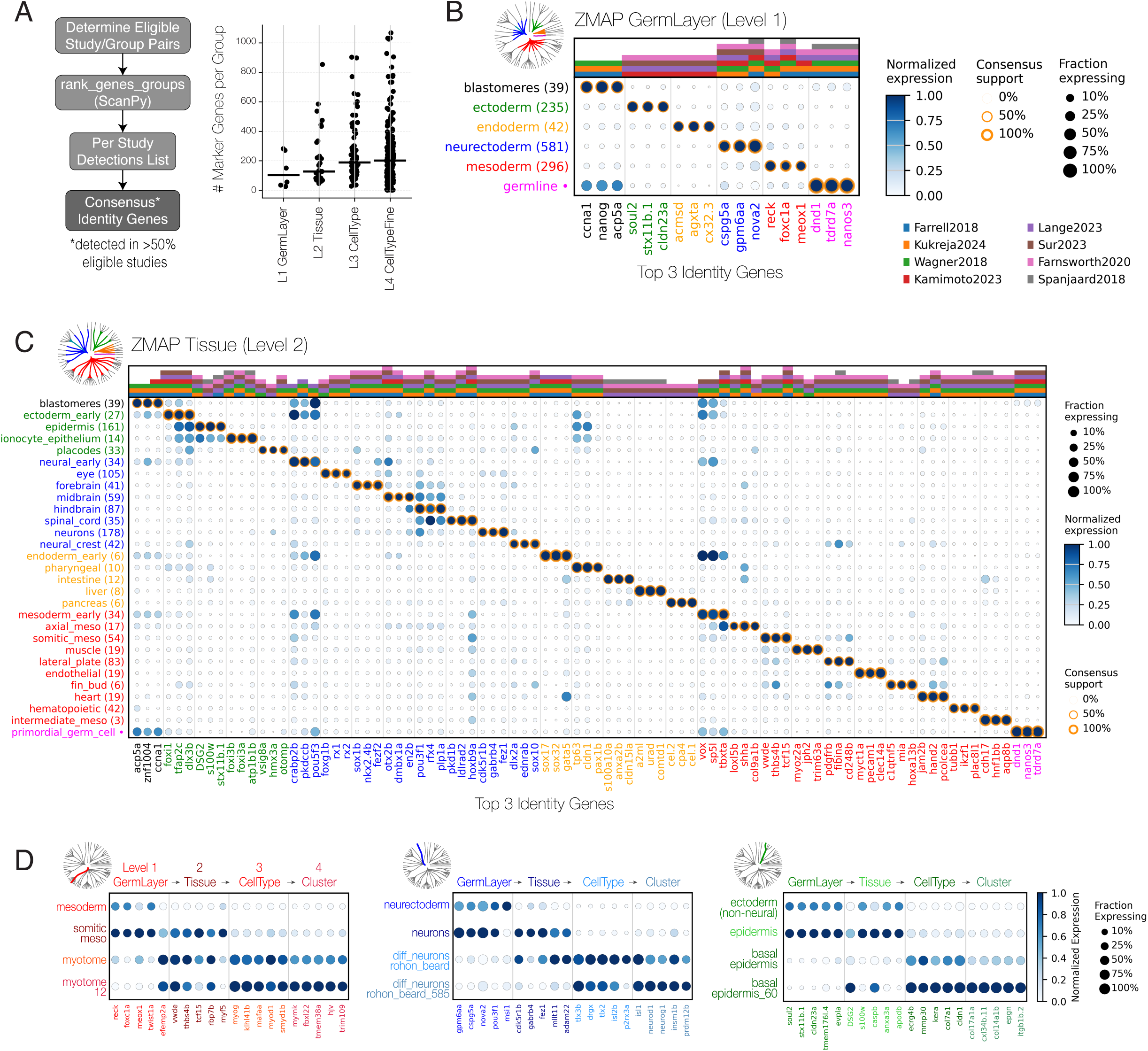
Consensus marker genes enable dissection of cell identity at multiple ontological levels. **(A)** Workflow for identification and ranking of consensus marker genes. Inset plots the number of marker genes per group across four levels of the ontology. **(B)** Dot plot showing consensus marker genes for GermLayer groups (ontology level 1). Inset radial tree (adapted from Fig. 2C) indicates ontology nodes represented in the dot plot. The top 3 ranked marker genes for each GermLayer group are shown. Dot color, size, and border ring indicate average expression level, fraction of expressing cells, and consensus support for each gene-group association, respectively. Stacked bars above the dot plot indicate individual studies in which each gene-group marker relationship was independently identified. Internal heterogeneity of each ontology node is indicated in parentheses as the number of constituent Cluster groups. Leaf nodes comprising a single Cluster group (n=1) are indicated as a dot. **(C)** Dot plot showing consensus marker genes for Tissue groups (ontology level 2). The top three ranked marker genes for each Tissue group are shown. Visual encodings are as in panel B. Stacked bars above the dot plot indicate the contributing studies supporting each gene-group marker relationship. **(D)** Dot plots showing consensus marker gene signatures along selected ontology paths: myotome cells (left), Rohon-Beard neurons (middle), and basal epidermis (right). Marker signatures exhibit progressive restriction at finer levels of the ontology hierarchy.

### Automated annotation using a Symphony-based ZMAP reference

To assess the suitability of ZMAP as a reference for automated annotation, we constructed a Symphony-based reference (*17*) following sub-sampling to minimize class imbalance (see Methods). Using this reference, query cells were then projected into the Harmony-corrected PCA space, ingested into the reference UMAP embedding, and annotated by distance-weighted k-nearest neighbors voting with Gaussian kernel weighting (Fig. 4A). Per-cell probability scores were used to filter low-confidence assignments (p ≥ 0.8).

**Figure 4.**
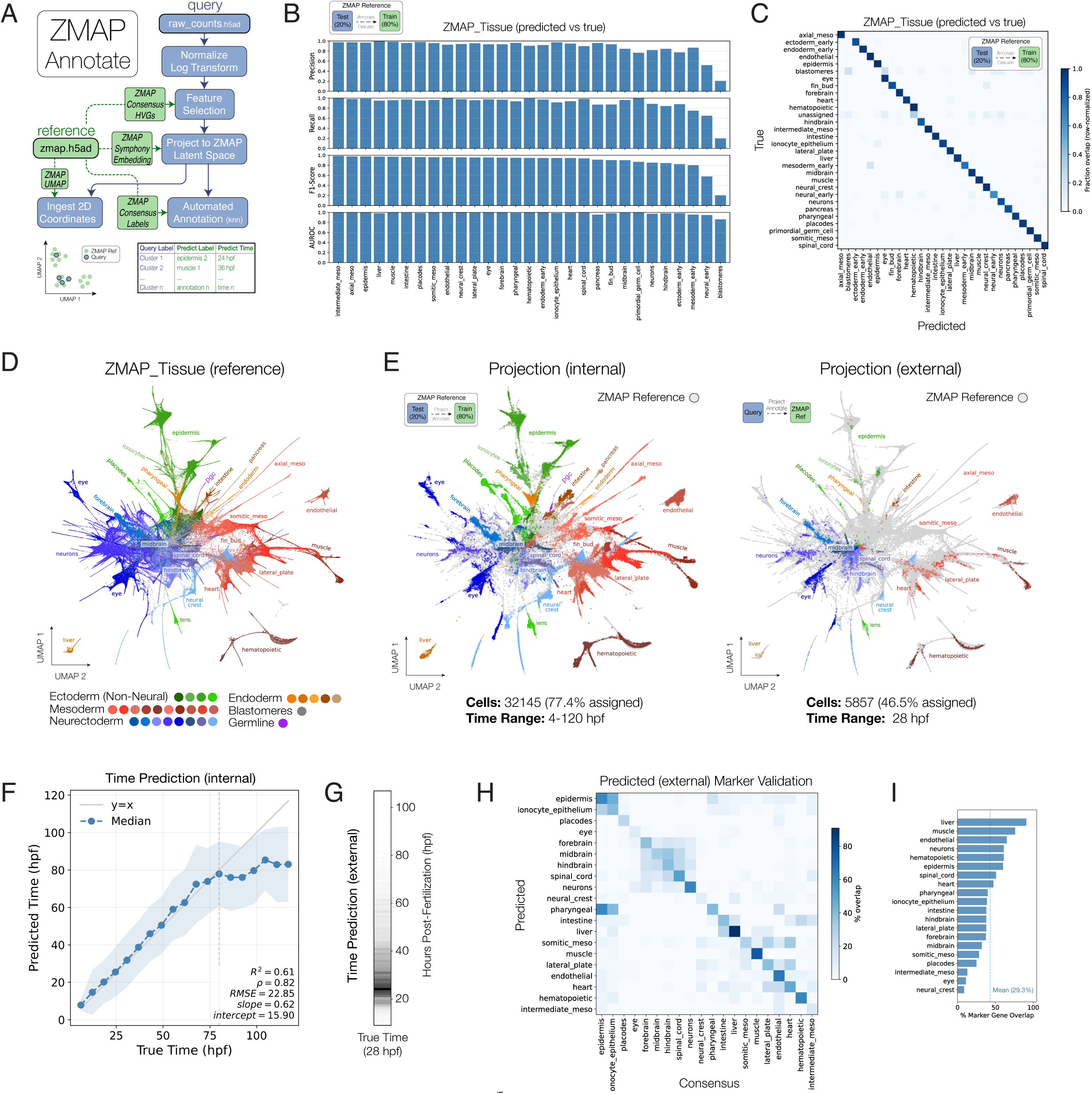
Automated annotation of zebrafish scRNAseq datasets. (A) Schematic of ZMAP annotation pipeline. Query cells are projected into the ZMAP embedding via Symphony and UMAP ingest, then annotated by distance-weighted kNN voting (see Methods). Performance was evaluated using both internal (20% holdout from the ZMAP reference) and external (non-ZMAP data) queries. (B) Per-label evaluation metrics: precision, recall, and F1 scores for kNN label transfer (internal query, ZMAP_Tissue level). (C) Confusion matrix of predicted vs. ground-truth tissue labels (internal query). (D) UMAP embedding of the ZMAP reference atlas; colors and labels indicate ZMAP_Tissue annotations. (E) Query cells projected into the reference UMAP. Reference cells are shown in grey; query cells are colored by predicted tissue label as in (D). Left, internal query (20% holdout); right, external query (non-ZMAP inDrops data). (F) Predicted vs. ground-truth developmental timepoint (internal query). Distance-weighted kNN time estimates show strong linear agreement across 3–72 hpf. (G) Density histogram of predicted timepoints (external query; collection timepoint 28 hpf) depicted as a heat strip. (H) Marker gene specificity matrix. For each predicted tissue group, the top 100 differentially expressed genes (Wilcoxon, log₂FC ≥ 1, q ≤ 0.05) were compared to curated ZMAP consensus marker gene lists for each reference tissue. Colors indicate % overlap. Strong diagonal signal confirms that predicted labels recover cell type-specific marker genes; off-diagonal entries reflect shared markers between related tissues. (I) Diagonal elements from (H): per-tissue marker overlap % between predicted DE genes and matched ZMAP consensus markers. Dashed line indicates mean marker gene overlap (29.3%) across all tissues.

We evaluated annotation performance using two independent query datasets: an internal holdout (20% of reference cells) and an external non-ZMAP inDrops scRNA-seq dataset. For the internal evaluation, 20% of reference cells were withheld, independently projected back into the reference embedding, and annotated by label transfer from the remaining 80% of reference cells. Following this procedure, each internal query cell carried: (1) its original “true” label, (2) a “predicted” label, and (3) a projected position in the reference UMAP. Per-label precision, recall, F1, and AUROC scores indicated strong agreement between predicted and ground-truth labels across most tissue types (Fig. 4B). Inspection of the corresponding confusion matrix revealed that while some misassignments were observed among “early” progenitor and “blastomere” labeled populations, labels corresponding to most other tissues were predicted with high accuracy (Fig. 4C).

To assess whether annotation performance could generalize beyond the ZMAP reference, we applied our pipeline to an independent inDrops scRNA-seq dataset (12,603 cells collected from ∼28 hpf embryos) not included in ZMAP construction. Both internal and these external query cells projected into biologically coherent regions of the reference UMAP (Fig. 4D-E), and recapitulated the distributions of reference tissues. Predicted developmental timepoints also showed strong linear correspondence with ground-truth time labels across the 3–72 hpf range for internal query cells (Fig. 4F). For external query cells, the distribution of predicted time labels was consistent with the experimental collection timepoint (28 hpf; see Fig. 4G). Together these data indicated that both cell type and temporal information were reliably encoded in the integrated embedding and could be recovered during projection.

We further validated external query annotations by comparing de novo differential expression signatures to curated ZMAP consensus marker gene lists. For each predicted tissue group, the top 100 differentially expressed genes were compared against ZMAP reference markers for all tissue types, yielding a marker specificity matrix (Fig. 4H). Notable diagonal signals confirmed that predicted labels recovered cell type-specific marker genes, while off-diagonal signals generally reflected marker sharing between related tissues. Per-tissue overlap percentage scores (Fig. 4I) demonstrated that the majority of predicted groups recovered substantial fractions of their corresponding reference markers, with a mean overlap of 29.3%.

Together, these results indicate that ZMAP functions as a reliable reference for automated annotation of new zebrafish scRNA-seq datasets, with performance validated by both classification metrics and marker gene recovery. This annotation pipeline is available as part of the <u>zmap-tools</u> Python package (PyPI), with full documentation and tutorials available at: https://zmap-tools.readthedocs.io.

### Interactive exploration of the ZMAP atlas

To facilitate ZMAP usage, we developed a web portal (https://wagnerlabucsf.github.io/zmap/) that facilitates interactive exploration of the atlas across genes, cell groups, and developmental time (Fig. 5). The main projection window displays either 2D or 3D versions of the UMAP embedding and supports interactive panning, zooming, filtering, and hover-based inspection of cell-level metadata (Fig. 5A). Dynamic overlay of selected numeric (e.g., gene expression, QC scores) and categorical (e.g. annotation) cell tracks is enabled by either text search or pulldown colormap selectors. The ZMAP portal supports both gene-centric and annotation-centric exploration. Gene/track text queries trigger lazy loading of cell data and dynamic updating of the embedding colormap, and simultaneously expose dot plots summarizing expression across annotation groups, timepoints, and studies (Fig. 5B). Users can similarly search and select annotation groups, filter clusters, or navigate the ZMAP ontology to highlight specific tissues or cell types and visualize group-level marker gene summaries (Fig. 5C). Additional interactive components support exploratory analyses such as: lasso selection of arbitrary cell groups within the embedding to examine enriched consensus identity genes, filtering of cell groups using interactive legends, and filtering of cells by developmental time using an interactive histogram.

**Figure 5.**
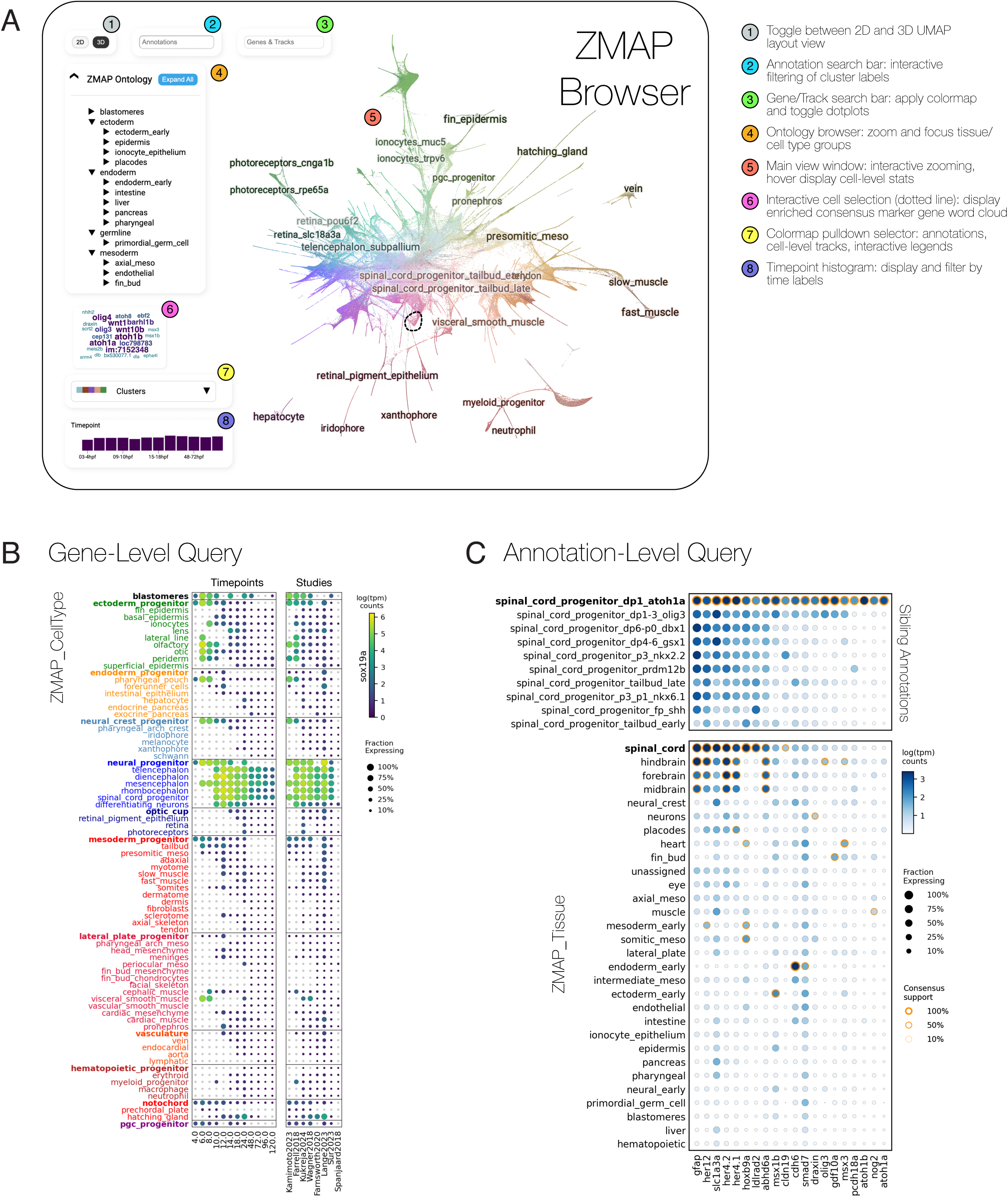
Interactive exploration of the ZMAP atlas via a web-based portal. (A) Overview of the ZMAP interactive web portal showing the primary 2D projection view and associated interface components. The projection window supports interactive panning, zooming, and hover-based inspection of cell-level metadata, with controls for toggling between 2D and 3D views, selecting color maps, filtering by developmental time, and browsing hierarchical annotations. (B) Example dotplot output from a gene/track level query. Gene selection triggers colormap updates to the embedding and opens a dot plot summarizing expression across annotation groups, time, and studies. (C) Example output from an annotation-centric query. Selecting a cell group through the ontology browser focuses the corresponding cells in the embedding window and opens a dot plot summarizing group-level identity marker gene expression. Top, view comparing marker gene expression among sibling groups. Bottom, view comparing amongst all Tissue groups.

### Conclusion

ZMAP provides the first unified, hierarchically annotated reference atlas that integrates diverse zebrafish scRNA-seq datasets into a coherent, harmonized framework. Several practical benefits derive from this large-scale cross-study integration. First, comparison of study-derived cell type labels revealed striking semantic concordance, which we interpret as reflecting shared underlying patterns of cell identity. These semantically related groups subsequently served as the basis for ZMAP annotations. Second, the cross-study nature of the atlas enables explicit separation of biological signal from technical variation. We leveraged this property to derive consensus identity programs for all nodes of the ZMAP annotation ontology.

The combination of a structured annotation ontology, a lightweight label transfer tool, and highly interactive visualizations distinguishes ZMAP from prior zebrafish scRNA-seq interfaces. ZMAP enables a wide range of potential downstream applications, including cell type annotation for new single-cell and spatial transcriptomics datasets, quantification of phenotypes affecting developmental tempo and heterochrony, and the identification of novel, ectopic, or pathogenic cell states. Future extensions of ZMAP will incorporate additional reference datasets and developmental stages. Moreover, we envision broad utility through the addition of modular data “tracks” representing perturbations, mutant phenotypes, barcoded lineage behaviors, spatial or anatomical metadata, and other modalities. Such tracks, summarized as vectors projected into the ZMAP reference embedding, will confer genome-browser-like modularity and extensibility. Together, these features position ZMAP as a foundational reference for quantitative analysis of zebrafish embryogenesis, and for the systematic integration of additional single-cell datasets into a common analytic framework.

## Materials and Methods

### Data Acquisition and FASTQ Preparation

Single cell RNA sequencing data were aggregated from 8 previously published zebrafish studies, Spanjaard et al, 2018 (*6*) (GEO: GSE81533, BioProject: PRJNA321866), Farrell et al 2018 (*3*) (GEO: GSE106474, BioProject: PRJNA417290), Wagner et al 2018 (*1*) (GEO: GSE112294, BioProject: PRJNA445487), Farnsworth et al 2020 (*5*) (BioProject: PRJNA564810), Kamimoto et al 2023 (*4*) (GEO: GSE145298, BioProject: PRJNA606682), Kukreja et al 2024 (*2*) (GEO: GSE269784, BioProject: PRJNA1123686), Lange et al 2023 (*8*) (BioProject: PRJNA940501), and Sur et al 2023 (*7*) (GEO: GSE223922, BioProject: PRJNA929041). To maximize comparability across studies and ensure sufficient transcriptome complexity for cross-dataset integration, we prioritized high-quality droplet-based scRNA-seq datasets with a minimum mean detection of >2,500 UMIs per cell. Sequencing data were obtained from NCBI Sequence Read Archive (SRA), Gene Expression Omnibus (GEO), associated Google Cloud Platform (GCP) buckets, Amazon Web Services (AWS) mirrors, and author hosted repositories, depending on availability. Datasets deposited as FASTQ files were downloaded directly. For datasets deposited only as aligned BAM files, FASTQ files were reconstructed as follows: cDNA reads were extracted using bedtools and bamtofastq, while cell barcode and UMI sequences were recovered from BAM tags and written to synthetic barcode FASTQ files with uniform base quality scores. Additional preprocessing steps were applied as required by library structure. 10X Chromium v1 libraries encoded cell barcodes and UMIs on separate reads, which were concatenated into a single barcode FASTQ prior to alignment. inDrops-based datasets required merging multiple barcode reads into a single barcode FASTQ. Within libraries, pooled or multi-lane sequencing runs were concatenated across lanes. Where necessary, short or empty reads were removed using cutadapt to enforce minimum length thresholds for both barcode and cDNA reads.

### Barcode and UMI Configuration

Barcode and UMI parsing parameters were set explicitly in STARsolo to reflect the sequencing chemistry used in each study. For 10X Chromium v1 libraries from Spanjaard2018, STARsolo was run with –soloType CB_UMI_Simple, --soloCBlen 14, --soloUMIstart 15, --soloUMIlen 10, and –soloBarcodeReadLength 0 (this setting permits variable read lengths), using merged barcode and UMI FASTQs. For 10X Chromium v2 libraries from Spanjaard2018 and Farnsworth2020, STARsolo was run with –soloType CB_UMI_Simple, --soloCBlen 16, --soloUMIstart 17, --soloUMIlen 10, and –soloBarcodeReadLength 0, with the barcode read provided as R2. For 10X Chromium v3 libraries from Kamimoto2023, Lange2023, and Sur2023, STARsolo was run with –soloType CB_UMI_Simple, --soloCBlen 16, --soloUMIstart 17, --soloUMIlen 12, and –soloBarcodeReadLength 0. For Drop seq libraries from Farrell2018, STARsolo was run with –soloType CB_UMI_Simple, --soloCBlen 12, --soloUMIstart 13, --soloUMIlen 8, and –soloBarcodeReadLength 1, using barcode and UMI sequences reconstructed from BAM tags. For inDrops v3 libraries from Wagner2018 and Kukreja2024, STARsolo was run with –soloType CB_UMI_Simple, --soloCBlen 16, --soloUMIstart 17, --soloUMIlen 6, and –soloBarcodeReadLength 1, using merged R2R4 barcode FASTQs.

### Cell Barcode Whitelists

Cell barcode whitelists were selected or constructed to match the sequencing chemistry of each dataset. The whitelist 10Xv1_737K-april-2014_rc.txt was used for Spanjaard2018 10X Chromium v1 libraries. The whitelist 10Xv2_737K-august-2016.txt was used for Spanjaard2018 10X Chromium v2 libraries and Farnsworth2020. The whitelist 10Xv3_3M-february-2018.txt was used for Kamimoto2023, Lange2023, and Sur2023. The whitelist inDrops_v3_whitelist.txt was used for Wagner2018 and Kukreja2024. For Farrell2018, a custom empirical whitelist was generated from published cell barcode lists. For Sur2023, additional library specific whitelists were generated directly from published barcode lists to ensure correspondence between reprocessed data and deposited count matrices. In all cases, whitelist matching was performed using –soloCBmatchWLtype 1MM.

### Reference Genome and Annotation

All datasets were aligned to a common zebrafish reference genome based on GRCz11 using the Lawson Lab v4.3.2 GTF file (*18*). Reference transcriptome and gene annotations were extended to comprise common transgenic sequences: mCherry2, mNeonGreen, mScarlet, mTagBFP2, kikGR1, mEOS4b, miRFP670, EGFP, sfGFP, Cas9, CRE, ERT2, Gal4ff, Tol2.

### Alignment and Quantification with STARsolo

All datasets were aligned using STAR version 2.7.10b in STARsolo mode. Across all studies, STARsolo was run in two pass alignment mode using –twopassMode Basic, with permissive run directory permissions (--runDirPerm All_RWX), decompression via zcat (--readFilesCommand zcat), and uniform handling of multimapping reads (--soloMultiMappers Uniform). For all datasets, STARsolo was configured to quantify Gene, GeneFull, and Velocyto features (--soloFeatures Gene GeneFull Velocyto). Barcode and UMI parsing parameters were set as described above for each technology. Cell calling was performed using EmptyDrops based filtering (--soloCellFilter EmptyDrops_CR) and/or by manual determination of UMI count thresholds using a weighted histogram (see below: Quality Filtering).

### Single-Cell Quality Filtering

Reads associated with individual cell barcodes were filtered using a multi-step quality control procedure applied to each of the 343 individual scRNAseq libraries included in this study: (1) Barcodes associated with low-quality cells were removed based on a library specific minimum UMI threshold estimated from a weighted transcript count histogram to distinguish encapsulated cells from droplets containing background RNA. (2) Barcodes with elevated mitochondrial transcript content (>5%) were excluded to remove stressed or dying cells. (3) Putative doublets were identified using Scrublet (*19*) and removed using score threshold (>0.20). Quality-filtered libraries were concatenated into a single AnnData object (“ZMAP_raw.h5ad”) containing library-, study-, and timepoint-level metadata and raw count matrices, and saved for downstream analysis.

### Normalization, Feature Selection and Dimensionality Reduction

Single-cell RNA-seq count matrices were processed using Scanpy (*20*), Palantir (*13*), and UMAP (*12*). Raw counts were retained prior to transformation. Cells were normalized to total counts (TPM; target sum = 1e6) and log transformed. Highly variable genes (HVGs, n=2411) were identified in a batch-aware manner (see below: Batch Correction). Gene expression values were variance-scaled, mean-centered, and identified HVGs were used for downstream dimensionality reduction. The number of informative principal components was estimated (n=291) using a permutation-based approach in which observed PCA eigenvalues were compared to eigenvalues obtained from PCA of randomized data (*1*, *21*). Principal components with eigenvalues exceeding the 95^th^ percentile of the randomized distribution were retained.

Low-dimensional structure was further characterized using diffusion maps and multiscale diffusion components computed with Palantir. Cell neighborhood graphs were constructed from the X_pca_harmony representation; leiden clustering and graph abstraction were used to characterize connectivity structure and to initialize visualization. UMAP embeddings were generated using the umap-learn API to enable additional parameter control, with settings chosen to emphasize global structure (n_neighbors = 500, min_dist = 0.4, spread = 5, repulsion_strength = 0.05) and initialized using PAGA-derived 2D coordinates. Embeddings were optimized for 5,000 epochs using Euclidean distance.

### Batch Correction

Batch-aware feature selection and integration were performed using Scanpy and Harmony (*11*), respectively. Highly variable genes (n=2411) were identified using sc.pp.highly_variable_genes with batch_key = ‘library_id’, prioritizing features that were consistently variable across biological replicates, studies, and collection timepoints. Batch effects were further corrected in PCA space using the Symphony (*17*) implementation of Harmony (sp.pp.harmony_integrate, key = ‘library_id’), with remaining parameters set to default. Convergence was reached after 2 iterations. Harmony-corrected components were used for downstream analyses requiring an integrated representation (e.g., neighborhood graph construction, UMAP visualization, community detection). Batch correction performance was assessed using scIB-metrics (*14*). To minimize confounding sources of variation, batch integration was assessed using a downsampled subset of 10,000 cells from five studies (Wagner 2018, Kukreja 2024, Lange 2023, Sur 2023, Farnsworth 2020) at a common developmental timepoint (24 hpf). Batch mixing was evaluated using integration Local Inverse Simpson’s Index (*11*, *14*) (iLISI) and the kNN Batch Effect Test (*15*) (kBET). Preservation of biological structure was assessed using graph connectivity and principal component regression (PCR) comparison, and overall batch removal was summarized using the Batch Removal Assessment Score (BRAS). Scores for each metric were compared between (i) a PCA embedding generated using the ScanPy default HVG selection method (sc.pp.highly_variable_genes with no batch key), (ii) a PCA embedding generated using batch-aware HVG selection, and (iii) a Harmony-corrected version of (ii).

### Clustering, Consensus Annotations & Ontology

Cell state neighborhoods were constructed from the Harmony-integrated low-dimensional representation using a kNN graph, and graph-based community detection was performed using the Leiden clustering algorithm (*16*). High-resolution Leiden clustering (resolution = 100; 1,506 resulting clusters) was used to define the primary units for annotation. Curated CellType labels were annotated manually by jointly considering semantic concordance among study-specific labels across integrated datasets and annotation resolutions, together with marker gene expression. Semantic relationships are formalized in a lookup table of study-specific labels associated with each curated CellType (Supplemental Table 2), where “score” quantifies the cumulative within-cluster fraction (among cells from that study) assigned to a given label, summed across all Leiden100 clusters within each CellType. Lower-resolution Leiden clusterings (e.g., Leiden resolutions 1, 5, 20), cluster connectivity, and marker gene expression were subsequently used to organize annotated clusters into a hierarchical annotation ontology. The final ZMAP ontology spans five levels: ZMAP-GermLayer, Tissue, CellType (primary draft annotation), CellTypeFine (manually refined cell subtypes), and Cluster (high-resolution Leiden clusters).

To construct the cross-study annotation ontology tree (Fig. 2C), relationships between annotation labels were inferred from the cell-cell kNN graph derived from the integrated embedding. kNN graph edges connecting cells assigned to different labels were counted and used to identify annotations that may not co-occur within the same cells but were locally adjacent in the integrated neighborhood graph. Label connectivities were computed across ZMAP annotation layers (ZMAP-GermLayer, Tissue, CellType) and selected study-specific annotation sets: Sur23_cluster.identity.super, Wagner18-CellTypeName, Kukreja24_CellState, Lange23_ontology_class, and Farnsworth20_CellType. A directed hierarchical graph was then constructed using networkx by linking labels across successive ontology levels. The resulting four-layer graph was rendered as a radial tree using D3.js.

### Pseudotime Calculations

Four complementary measures of developmental progression were computed per cell. (1) kNN-smoothed time: experimental collection timepoints (time_id) were rank-transformed and smoothed over the cell-cell kNN graph (adata.obsp[‘connectivities’]). For each cell, the mean time_id of its k nearest neighbors was assigned, iterated for three rounds, and rescaled to the unit interval (0–1). (2) Palantir pseudotime: Palantir (*13*) was run on the first 100 min-max-scaled multiscale diffusion components. A single start cell was randomly selected from the earliest timepoint (fixed seed) and included with max-min-sampled waypoints and diffusion boundary cells. (3) Diffusion pseudotime (*22*) (DPT): Using the same start cell as Palantir, diffusion maps (100 components) and DPT were computed with sc.tl.dpt. (4) CytoTRACE: CytoTRACE (*23*) was computed using CellRank2 (*24*) from TPM-normalized, log-transformed expression values derived from raw counts.

### Derivation of Consensus Identity Genes

Consensus identity genes were defined as genes that consistently distinguished a given cell group across multiple studies after dataset integration. The procedure consisted of three stages: (1) per-study marker detection, (2) cross-study aggregation, and (3) unified ranking of genes based on complementary criteria.

### Per-study detection

For each annotated group (e.g., GermLayer, Tissue, or CellType) within each study, one-versus-rest differential expression was performed using the Wilcoxon rank-sum test on log-transformed TPM values using sc.tl.rank_genes_groups (*20*). Both target and reference groups were subsampled to 500 cells each. A study-group pair was considered eligible if it comprised at least 10 cells and accounted for at least 1% of the group’s total cell count in the integrated dataset. This filter prevented very small or under-represented study-group pairs from contributing spurious detections to the consensus. Within each eligible study-group pair, a gene was considered detected if it met all of the following criteria: adjusted p-value ≤ 0.01, log₂ fold change ≥ 1, expression detected in at least 10% of group cells, and a ≥ 2-fold enrichment of expressing-cell frequency relative to all other groups.

### Cross-study aggregation and consensus support

For each gene within each group, all studies that independently detected the gene were recorded. The number of supporting studies (support) was divided by the number of studies eligible for that group to yield a consensus support ratio. Each gene was then assigned a median rank across supporting studies, providing a within-group ordering of marker strength. To provide global context, all candidate genes identified in any study were subsequently re-tested across the full integrated atlas using a one-versus-rest differential expression test (sc.tl.rank_genes_groups), yielding global log₂ fold changes, adjusted p-values, and test statistics. For each gene and group, mean and median in-group and out-group expression fractions were also recorded.

### Composite ranking

To prioritize genes that were both statistically robust and biologically distinct, we computed a hierarchy of rank-based summary scores within each group. “<u>Contrast Rank</u>” captured “classic” differential expression strength by combining the within-group ranks of adjusted p-value and log₂ fold change. “<u>Specificity Rank</u>” summarized cell-type distinctiveness using the ranks of enrichment and marker gene recurrence, prioritizing genes highly enriched within a group and infrequently reused elsewhere. “<u>Prevalence Rank</u>” reflected the mean fraction of cells expressing the gene within the group. “<u>Consensus Rank</u>” captured reproducibility across studies using the support ratio. Finally, for each gene (within each group), an “<u>Overall Rank</u>” was determined by averaging across all individual rank scores, producing a single composite measure that balanced effect size, specificity, and reproducibility. Consensus Identity Genes, their ranks and associated statistics are summarized in Supplemental Table 3.

### Automated Annotation

To facilitate automated annotation of query scRNAseq datasets, a Symphony reference h5ad object (*17*) was constructed from a curated subset of annotated ZMAP cells. To maintain balanced representation across developmental stages and cell types, cells were subsampled by both time window and cell identity (CellType); within each cell type-time intersection, up to 400 cells were retained, or all available cells if fewer were present. Harmony-adjusted embeddings (X_pca_harmony) and batch correction factors were subsetted to match the retained cells, and neighborhood graphs were recomputed using 100 neighbors. The resulting data subset consisted of 207,755 cells. Query data were preprocessed from raw counts by library-size normalization to a target sum of 10^6^ followed by log normalization (log-TPM counts). Queries were mapped to the ZMAP reference using Symphony, first by projecting into Harmony-adjusted PCA space (X_pca_harmony) using sp.tl.map_embedding, then ingesting into the reference UMAP with sp.tl.ingest. Labels were transferred with a distance-weighted k-nearest neighbors procedure operating in the Harmony-corrected PCA space using k=25 and cosine distance.

Neighbor votes were weighted by a Gaussian kernel (σ = per-cell median neighbor distance). Quality control thresholds were applied to accept predictions (p ≥ 0.8). Predicted groups with small numbers of assigned cells (n<15) were excluded. Developmental timepoints were predicted for each query cell by computing a distance-weighted trimmed mean of the reference neighbor time values, using the same Gaussian kernel weights applied during label voting.

When ground truth was available (e.g. during test/train split analyses), accuracy, macro-precision/recall/F1, per-class AUROC, and confusion matrices were computed on accepted predictions. For visualization, query cells with predicted labels and ingested coordinates were overlaid on the Symphony reference UMAP.

### Single-Cell Dissociations

Zebrafish embryos were transferred to a 1.5 mL Protein LoBind Tubes (Fisher Scientific, 13698794) prior to dissociation. 29 hpf embryos were enzymatically dissociated using a prewarmed Dissociation Mix (30°C) composed of 460 µL of Trypsin-EDTA (0.25%) (Gibco, Cat# 25200-056), 40 µL of 100 mg/mL Collagenase/Dispase (Sigma-Aldrich, Cat# 10269638001), 40 µL of Dnase I (1 mg/mL) (StemCell Technologies, Cat# 07900). Danieau Buffer was removed, leaving ∼10 µL of buffer, and 150 µL Dissociation Mix was added to dechorionated embryos using a P200 pipette. Embryo tissue was mechanically dissociated by pipetting for 30 seconds, followed by a 2 mins incubation at 30°C. 390 μL Dissociation Mix was added to the sample and mixed for 30 seconds by pipetting. For 10 minutes, samples were incubated at 30°C and mixed for 30 seconds at 2-minute intervals by pipetting, to dissolve remaining tissue clumps. 800 µL DMEM supplemented with 10% FBS (Thermo Fisher Scientific, Cat# SH30071.03HI) equilibrated at 30°C was used to neutralize the samples. Samples were filtered through a 40-µm cell strainer (Fisher Scientific, Cat# 0877123) before centrifuged in a swinging bucket rotor at 700 rcf for 5 minutes at 4°C. The supernatant was removed, and cells were resuspended in 200 µL of a Wash Buffer composed of 1x PBS and 1% BSA (Fisher Scientific, 501215315), and resuspended in an additional 200 µL of Wash Buffer for a total volume of 400 µL. Samples were washed twice before resuspending in 400 µL of Resuspension Buffer (10% OptiPrep (Sigma, D1556), 1x PBS, and 0.1% BSA in ddH2O) for single-cell microfluidic barcoding.

### inDrops Single-Cell Barcoding

An ONYX microfluidics platform (Droplet Genomics, DG-ONYX), V2 barcoding PDMS chips (Droplet Genomics DG-CBC2-80-10), and inDrops V3 hydrogel beads (*25*) were used to encapsulate single cells for cDNA barcoding. Hydrogel beads were centrifuged, washed, and loaded into the microfluidic setup using a syringe and PTFE tubing (inlet 1) prior to cell dissociations; beads were kept out of direct light during setup. Reverse transcription (RT) master mix (chilled; inlet 2), Droplet Stabilizing Oil (DG-DSO-15, inlet 3), and dissociated cell suspensions (inlet 4) were each prepared and similarly loaded onto the ONYX instrument. Flow rates were adjusted to obtain ∼1 nanoliter-sized droplets and a hydrogel bead occupancy of >90%. Post-encapsulation, emulsions were collected and exposed for 7 minutes to 365 nm UV light to release primers. The emulsion was split into ∼2000 cell equivalent volumes and aliquoted into 0.2 mL PCR tubes containing 30 μL Droplet Stabilizing Oil and 200 μL mineral oil. Within-droplet template-switching reverse transcription (Maxima H Minus RT; Thermo EP0751) reactions were then performed on a PCR block at 42 °C for 60 mins, followed by 5 mins at 85 °C, and held at 4 °C. Mineral and droplet stabilizing oil were then removed, and 2 uL of EDTA and 50 μL of Emulsion Breaker (Droplet Genomics DG-EB-1) were added. Barcoded cDNA samples were vortexed, briefly spun, transferred into a Zymo Spin Column (VWR, 76004-136), spun at 16000g for 1 min, collected into a clean 1.5mL tube, and chased with 50 μL of dH2O before storage at -80 °C.

### Preparation and Sequencing of inDrops cDNA Libraries

Barcoded purified cDNA samples were equilibrated to room temperature for two rounds of 0.8x SPRI bead-based size selection (AMPure XP Beads; Beckman A63881). Terra Direct PCR master mix (Takara 639270) and universal primers were added to each library, and cDNA was PCR-amplified followed by two 0.6x SPRI purification steps. Yield and size distributions of amplified cDNA libraries were assessed using an Agilent 2200 TapeStation system (G2964AA) with a D5000 ScreenTape (5067–5588) and D5000 Reagents (5067–5589). Library fragmentation was performed using NEBNext Ultra II FS reagents (New England Biolabs).

Enzymatic fragmentation reactions were prepared on ice, vortexed, spun down, and cDNA fragmentation and simultaneous dA-tailing were performed on a thermocycler, followed by a 0.6-0.8x double-size SPRI selection. Adapter ligation mix was incubated in the thermocycler for 15 mins at 20°C, followed by a 0.8x SPRI purification, a PCR indexing step, and another double 0.6-0.8x SPRI size selection. Final samples were quantified with the Qubit dsDNA BR Assay Kit (Q32850) and a second TapeStation analysis prior to sequencing. 8 biological replicate inDrops libraries were sequenced on NextSeq 2000 P2 FlowCell with cycle allocations as follows. Read 1: 16 cycles (Cell Barcode Part 1); Read 2: 65 cycles (cDNA); I5 Index Read: 8 cycles (Cell Barcode Part 2); I7 Index Read: 6 cycles (Sample Index). Sequencing data were mapped and processed as described above.

### Interactive Web Portal

The ZMAP interactive portal was implemented as a client-side web app for visualization of single-cell embeddings. The portal was built using DataMapPlot, which wraps deck.gl, D3.js and other javascript components, and extended with custom javascript functions to enable gene-and annotation-level queries linked to dotplot summary visualizations. Embedding coordinates, annotation labels, and cell-level metadata were exported using Python and serialized into static files for deployment. For each gene, log(tpm) expression values are cloud-hosted as compressed Apache arrow vector files, retrieved in response to user queries, and used to update the embedding colormap and associated legend. Annotation queries and ontology-based selection are used to identify and highlight subsets of cells and to generate group-level consensus marker gene summaries. Cell subsets can be selected directly within the embedding using a free-form lasso tool and used to compute enrichment of consensus identity marker genes, visualized as a word cloud. Additional interface elements include a colormap selector with synchronized legends, a developmental time histogram used to filter cells by time window. The portal is deployed as a static web application, and its components are hosted via a content delivery network (Cloudflare).

## Data and Code Availability

Python-based functions and an API are available at: https://github.com/wagnerlabucsf/zmap-tools and via pip: https://pypi.org/project/zmap-tools/. ZMAP references (raw and processed ScanPy AnnData objects), consensus marker tables, and trained classifier objects can be downloaded via the zmap-tools API, from NCBI-GEO, and/or via a web portal hosted at https://wagnerlabucsf.github.io/zmap/, which also provides access to interactive plots. Cell-level quality control filter steps used custom Python functions available at: https://github.com/wagnerde/scTools-py (release v2023.11.13).

## Acknowledgements

The authors thank J.Gagnon, J.Farrell, L.Solnica-Krezel, T.Tsai, C. Mosimann, and the entire Wagner Lab at UCSF for feedback on the ZMAP project and insightful discussions.

## Author Contributions

N.A.S. performed inDrops experiments and contributed to ZMAP atlas construction, dataset processing, and systematic testing and validation. Y.S. contributed to development of the ZMAP annotation tools, web portal, dataset processing and validation. D.E.W. conceived and supervised the study, performed the scRNA-seq data analysis, developed the ZMAP atlas and web portal. All authors contributed to manuscript preparation and approved the final manuscript.

## Supplementary Information

### Description of Additional Supplementary Files

**Supplemental Table 1: Cluster-level annotations.** Leiden cluster identities (resolution = 100; 1,506 clusters) and their assigned CellType labels, derived from majority vote across contributing study annotations. Top 20 consensus identity marker genes (overall rank) are also displayed for each cluster.

**Supplemental Table 2: Mapping of study labels.** For each ZMAP CellType annotation, this table reports the most frequent corresponding label from component studies (Wagner18, Kukreja24, Farnsworth20, Lange23, Sur23), along with its fractional representation, total cell count, and cell count for the top label, across all available annotation fields provided by each label set.

**Supplemental Table 3: Consensus identity genes.** Long-form table of consensus marker genes for each ZMAP ontology group across all annotation levels (GermLayer, Tissue, CellType, CellTypeFine), including differential expression statistics, effect sizes, expression prevalence, and cross-study reproducibility scores.

